# Computationally Optimized H1 and H3 Hemagglutinin Messenger RNA Vaccines Confer Broad Protective Immunity Against Modern Influenza Viruses

**DOI:** 10.64898/2026.09.04.749505

**Authors:** James D. Allen, Jessica M. Medina, Camila Caetano, Matthew H. Thomas, Amanda Lynch, Ted M. Ross

## Abstract

The hemagglutinin (HA) glycoprotein of seasonal influenza viruses undergoes continual antigenic drift, contributing to vaccine mismatch and reduced effectiveness of strain-specific seasonal vaccines. Although vaccination remains the most effective strategy for preventing influenza disease, conventional egg-based vaccine production requires several months and may not keep pace with viral evolution. Messenger RNA (mRNA) vaccines offer a promising alternative because they can be rapidly updated to encode emerging antigens and are manufactured through a scalable, cell-free process that avoids propagation associated adaptations. To address the challenges of antigenic drift and vaccine mismatch, we combined mRNA vaccine technology with Computationally Optimized Broadly Reactive Antigens (COBRA) to develop broadly protective influenza HA vaccines. These COBRA H1 and H3 mRNA HA vaccines elicited robust antigen-specific IgG, hemagglutination inhibition (HAI), and neutralizing antibody responses against diverse historical and contemporary influenza strains in cohorts of influenza naïve and pre-immune mice. Vaccination also induced strong cellular immunity, characterized by the expansion of antigen-specific antibody and cytokine secreting cells. These responses were further enhanced in animals with pre-existing influenza immunity, as demonstrated by an increased frequency of IFN-γ producing cells recognizing conserved HA stalk-based peptides. Together, these findings demonstrate that COBRA HA encoding mRNA vaccines can effectively leverage immunological memory while expanding responses to conserved HA epitopes, supporting improved protection against antigenically drifted strains. Thus, the combination of broadly reactive COBRA HA antigens with a rapidly adaptable and manufacturable mRNA platform represents a promising strategy for next-generation influenza vaccination.

**Importance:** Recent influenza seasons have demonstrated that vaccine mismatch can have significant public health consequences, leading to increased disease burden, hospitalizations, and transmission. Modern mRNA vaccines offer a promising alternative to traditional egg-based vaccines because they can be rapidly manufactured and updated to better match emerging viral variants while eliciting strong antiviral immune responses. Importantly, the flexibility of the mRNA platform enables the expression of custom-designed antigens optimized for broad protection. Advances in computational antigen design now make it possible to incorporate immunologically relevant epitopes from multiple circulating viruses into a single vaccine antigen. By expanding immune responses beyond strain-matched protection, these broadly reactive antigens can elicit antibodies that recognize both dominant and emerging viral variants. As a result, they have the potential to establish immune memory against antigenically diverse strains before they become prevalent in the population, improving vaccine effectiveness during periods of antigenic drift.

## Introduction

Influenza viruses cause acute respiratory disease characterized by fever, body aches, fatigue, and respiratory distress (1, 2). These viruses infect 5-15% of the global population annually, resulting in 290,000 to 650,000 deaths worldwide (3, 4). Although vaccination is an effective preventative strategy, influenza is an antigenically variable virus that undergoes continual antigenic drift to evade host immune pressure (5, 6). This process is driven by an error prone viral RNA-dependent RNA polymerase which introduces amino acid substitutions in the hemagglutinin (HA) and neuraminidase (NA) glycoproteins, thereby enabling escape from preexisting immunity (7–9). These substitutions are essential to the survival of the virus and complicate predictions of viral evolution (10). Currently, seasonal influenza vaccines are reformulated annually based on the strains that are predicted to predominate in the upcoming year (11, 12). However, current vaccines, typically composed of inactivated virus or recombinant HA, are largely strain-specific and require three to six months to manufacture at scale (13, 14). During this interval, the continual antigenic drift of circulating viruses can result in vaccine strain mismatch and reduced effectiveness (11, 15). Therefore, strategies that broaden antigen-derived protective immunity while accelerating vaccine production are urgently needed.

Most commercially available inactivated influenza vaccines are produced in chicken eggs, whereby viral isolates are propagated and subsequently inactivated with beta-propiolactone (BPL) or formaldehyde to render the virus incapable of replication (16, 17). This manufacturing process is time consuming, dependent on a reliable egg supply, and can result in egg-adaptative mutations that may reduce vaccine efficacy (18–20). Although egg adaptation can be minimized by propagating viruses in mammalian cell cultures, chemical inactivation is still required prior to use, which can decrease antigenicity and vaccine effectiveness (16, 17, 21). Alternatively, recombinant HA subunit vaccines can be rapidly updated in response to emerging viral outbreaks, produced in mammalian cell cultures, and manufactured at scale within approximately three months (22–25). However, these vaccines are often poorly immunogenic when administered alone and typically require high antigen doses or adjuvants to elicit protective immune responses (26–28).

Messenger RNA (mRNA) vaccine platforms enable faster manufacturing than traditional egg-based vaccines (29). Notably, mRNA vaccines for SARS-CoV-2 advanced from sequence acquisition to phase 1 clinical trials in approximately two months (15, 30). Similar to recombinant vaccines, mRNA vaccines can be rapidly updated to encode the most relevant antigens in response to pathogen evolution, thereby reducing the risk of vaccine mismatch and potentially enhancing global response strategies to emerging influenza strains (31). This cell-free manufacturing process also enables flexible, scalable, and rapid production without the risk of viral adaptation. However, due to their high molecular weight and inherent chemical instability, mRNA vaccines are often encapsulated in lipid nanoparticle (LNP) carriers to protect the mRNA payload and facilitate cytoplasmic delivery (32, 33). Beyond their delivery function, LNPs also possess intrinsic adjuvant properties and contribute to the production of robust humoral and cellular immune responses (18, 34, 35).

In addition to manufacturing speed and immune stimulation, antigen selection plays a critical role in determining seasonal influenza vaccine efficacy. Wild-type vaccine antigens are typically strain-specific and therefore provide limited protection against antigenically drifted influenza strains (11). To address this limitation, an *in-silico* methodology was developed to generate Computationally Optimized Broadly Reactive Antigens (COBRAs) for influenza HA (36, 37). COBRA HA antigens are designed using contemporary wild-type HA sequences collected through global influenza surveillance networks, which are layered using a consensus-building approach (38). This strategy assigns comparable weight to both dominant and sub-dominant strains within each season, resulting in a final antigen that incorporates immunologically relevant epitopes from a diverse set of recently circulating viruses. COBRA HA antigens elicit broad protective immune responses in mice, ferrets, and non-human primates, and can be expressed across multiple vaccine platforms including recombinant protein, virus like particle (VLP), and mRNA-based vaccines (39–44). Importantly, COBRA HA antigens can be updated in near real time to reflect actively circulating viruses, thereby broadening vaccine-induced immune responses against clinically relevant strains (45, 46). Thus, combining COBRA HA antigens with a rapidly manufacturable vaccine platform like mRNA, may offer a highly advantageous strategy for the future of influenza vaccination.

In this study, cohorts of influenza naïve and pre-immune mice were vaccinated with mRNA LNPs encoding COBRA Y2 H1 and NG5 H3 HA antigens. The vaccine induced humoral and cellular immune responses were evaluated for immunologic breadth against panels of H1N1 and H3N2 vaccine strains from the past decade. A two-dose COBRA mRNA LNPs regimen elicited robust hemagglutination inhibition (HAI) and virus-neutralizing antibody responses, increased the frequency of HA-specific antibody secreting cells, and conferred protection against weight loss and clinical disease following challenge with a contemporary H1N1 virus.

## Results

### Influenza naïve and pre-immune mice vaccinated with COBRA mRNA LNP vaccines had protective antibody titers against H1N1 and H3N2 vaccine strains from the past decade

A cohort of influenza naïve and pre-immune female DBA/2J mice (n=112) was vaccinated intramuscularly in a prime-boost regimen with monovalent mRNA LNPs encoding either the Y2 H1COBRA HA, NG5 H3 COBRA HA, or a bivalent formulation containing both mRNAs (**Fig. 1**). Serum collected 14 days after the second vaccination (day 42) was analyzed for influenza-specific IgG responses (**Fig. 2**). Pooled sera from mice immunized with Y2 HA containing vaccines had higher concentrations of H1 HA-specific IgG antibodies than sera from mice immunized with NG5 HA containing or mock vaccines against the Y2, BR/18, and cH6/1 rHA proteins (**Fig. 2A-C**). Conversely, pooled sera from mice immunized with vaccines encoding the NG5 HA exhibited higher concentrations H3 HA-specific IgG antibodies compared with Y2 HA or mock vaccinated groups against the NG5, TS/20, and cH7/3 rHA proteins (**Fig. 2D-F**). At the individual level, both influenza naïve and pre-immune mice vaccinated with mRNA LNPs encoding Y2 or NG5 HA had significantly higher Y2 HA- or NG5 HA-specific total IgG endpoint titers in both monovalent and bivalent formulations compared with the mock vaccinated controls (p < 0.05) (**Supp. Fig. 1A, B**).

**Figure 1.**
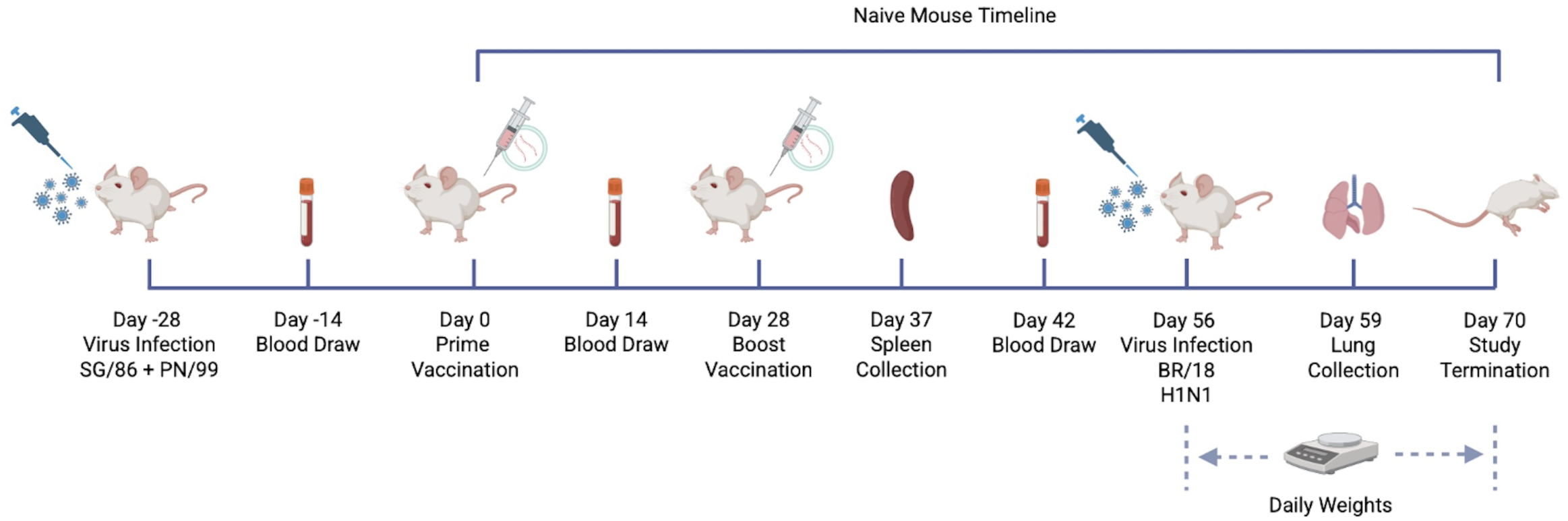
Mouse vaccination and challenge timeline. A cohort of 112 DBA/2J mice were divided into two groups. The first group (n=56) was intranasally infected with H1N1 (SG/86) and H3N2 (PN/99) on day −28 of the study. Four weeks later, on day 0, all mice in both groups (n=112) were intramuscularly vaccinated with 1 μg of mRNA LNPs encoding COBRA Y2 or NG5. The vaccines were administered in monovalent and bivalent formulations to groups of n=15 mice, and a mock group was established by vaccinating n=11 mice with PBS. All mice were boosted with the same vaccines on day 28 as they received on day 0. Blood was collected from each mouse on days −14, 14, and 42. Spleens were collected from 3 mice in each group on day 37. On day 56, the mice were intranasally infected with H1N1 virus (BR/18). Following infection mice were monitored for weight loss and clinical symptoms of infection for 14 consecutive days. Lungs were collected from n=3 mice in each group on day 59. The study was terminated on day 70. This figure was created using Biorender.com

**Figure 2.**
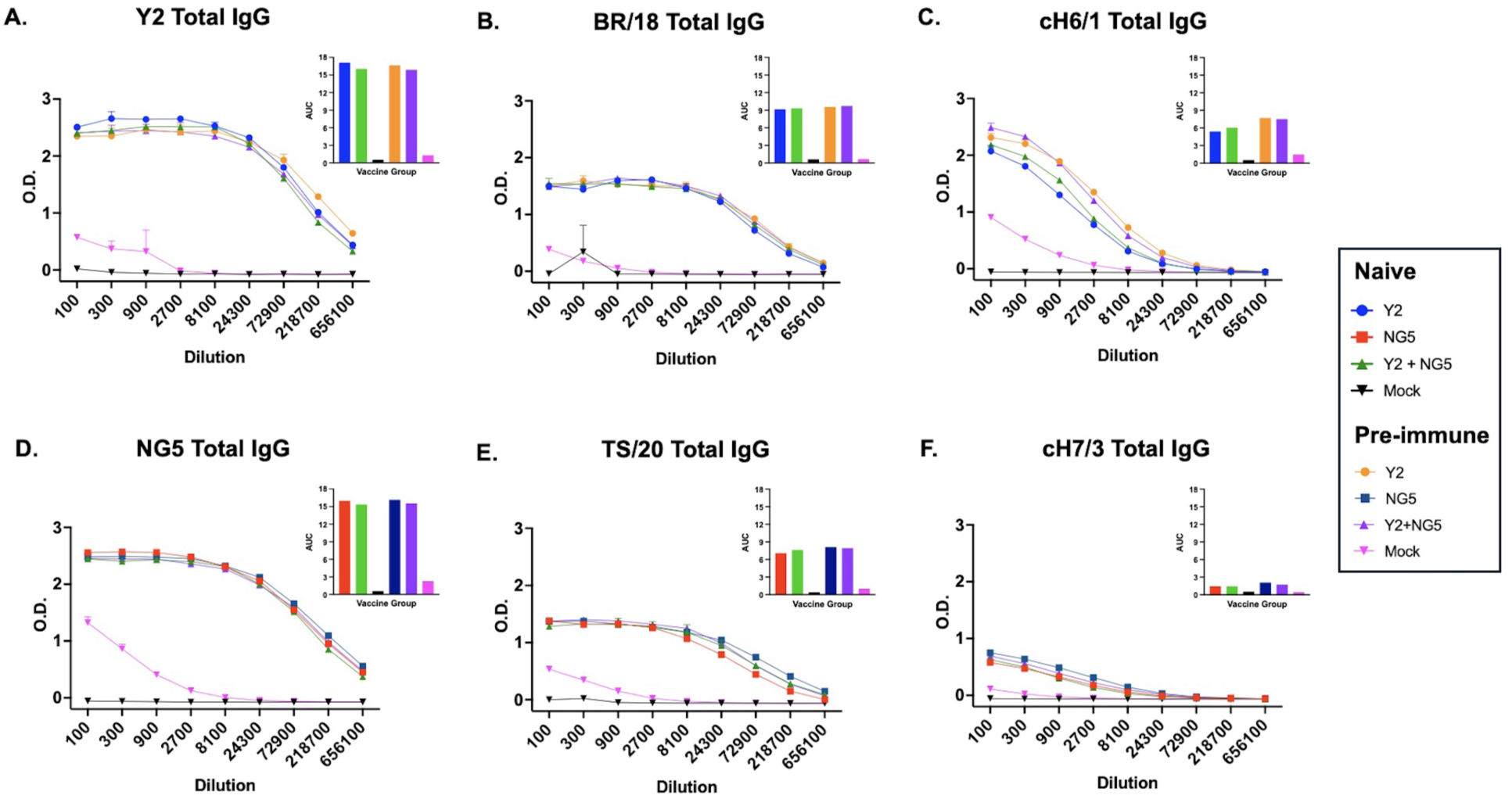
Pooled day 42 antigen specific total IgG titers. Serum collected following the second vaccination, day 42, was pooled for each group and assayed for total IgG content specific to Y2 (A), BR/18 (B), cH6/1 (C), NG5 (D), TS/20 (E), and cH7/3 rHA proteins by ELISA. Optical Density (O.D.) is plotted as a function of serum dilution, and the area under the curve (AUC) is reported in each panel. Naïve mice were vaccinated with mRNA LNPs encoding Y2 (blue), NG5 (red), Y2 + NG5 (green) HA proteins, or mock (PBS) (black). Pre-immune mice were vaccinated with mRNA LNPs encoding Y2 (orange), NG5 (navy blue), Y2 + NG5 (purple) HA proteins, or mock (PBS) (pink).

After a single immunization, all but one mouse vaccinated with Y2 HA mRNA developed sero-protective (≥1:40) hemagglutination inhibition (HAI) titers against BR/18, whereas only a few mice achieved protective titers against VC/22 (**Supp. Fig. 2A, B**). In comparison, following one immunization, naïve mice vaccinated with NG5 HA encoding mRNA did not generate sero-protective HAI titers against HK/19 or MA/22, whereas a subset of the pre-immune mice had sero-protective HAI titers against these viruses (**Supp. Fig. 2C, D**). Following the second immunization, all influenza naïve and pre-immune mice vaccinated with Y2 HA containing vaccines had sero-protective HAI titers against every historical H1N1 vaccine strain tested from 2009 to 2022 (**Fig. 3A, B**). Mice vaccinated with mock or monovalent NG5 HA had no detectable HAI activity against any virus in the H1N1 panel, with titers significantly lower than those elicited by the Y2 HA mRNA LNP vaccination (p < 0.01). No significant differences in HAI titers were observed between naïve and pre-immune mice or between monovalent and bivalent formulations containing the Y2 HA (**Fig. 3A, B**). For the H3N2 panel, influenza-naïve mice vaccinated with mRNA encoding NG5 achieved mean sero-protective HAI titers against five of the seven the historical strains tested (**Fig. 3C**). Pre-immune mice vaccinated with Y2 + NG5 also developed mean protective HAI titers against five of the seven H3N2 viruses, while mice receiving the monovalent NG5 HA formulation also exhibited mean protective HAI titers against the same strains and the DC/23 strain (**Fig. 3D**). In contrast, mice vaccinated monovalent Y2 HA or mock vaccines did not have detectable HAI titers against any H3N2 virus, with responses significantly lower than those elicited by NG5 HA containing vaccines (p < 0.01). No statistical differences in HAI titers were observed between monovalent and bivalent COBRA HA mRNA LNP formulations, however pre-immune mice vaccinated with the monovalent NG5 HA formulation had significantly higher HAI titers than naïve mice vaccinated with same formulation against the KS/17 and MA/22 strains (p < 0.05) (**Fig. 3C, D**).

**Figure 3.**
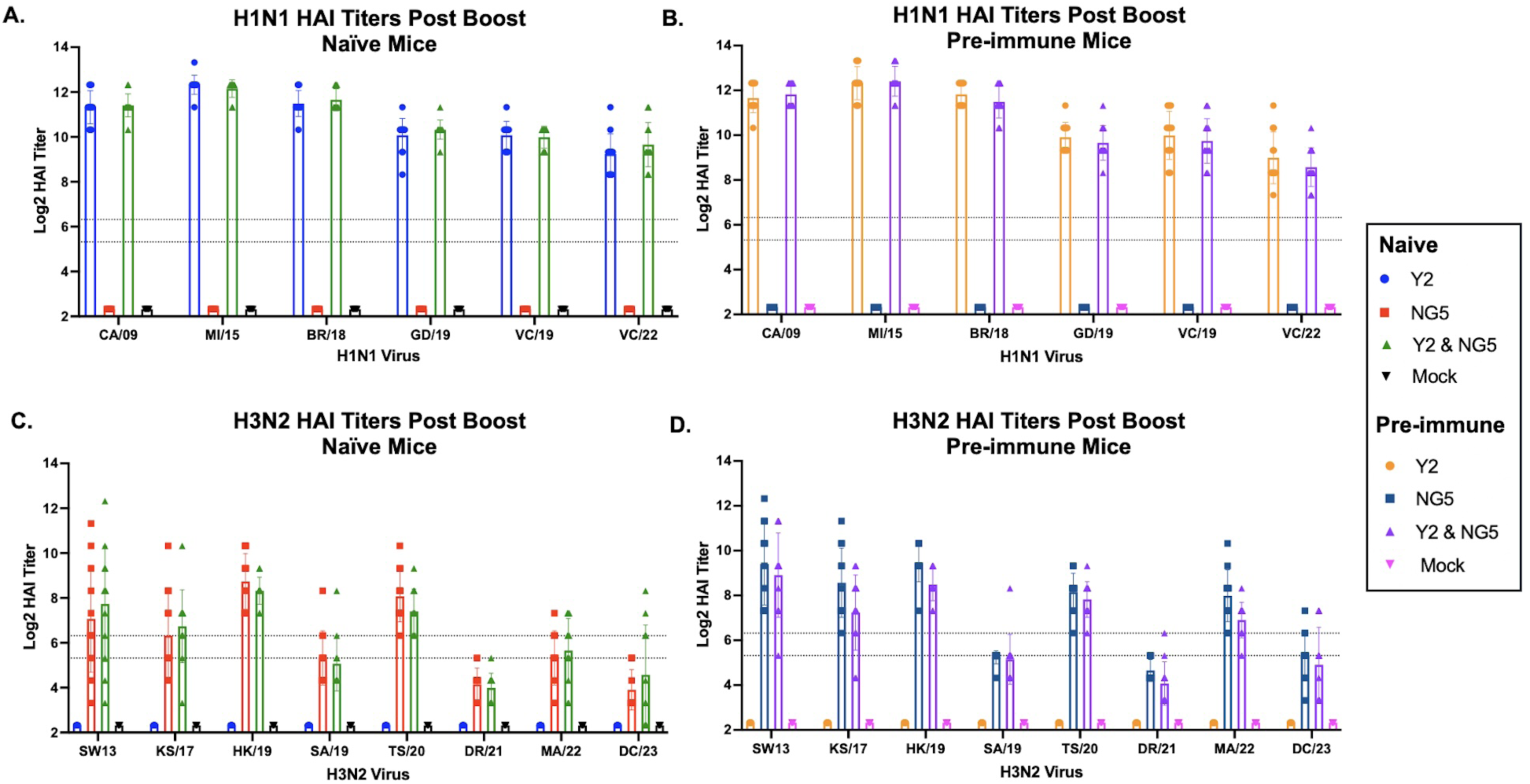
Day 42 HAI specific antibody response. Serum samples collected from naïve (A, C) and pre-immune mice (B, D) two weeks after the second vaccination, day 42, were assessed for HAI reactive antibodies against historical H1N1 (A, B) and H3N2 (C, D) vaccine strains. Naïve mice were vaccinated with mRNA LNPs encoding Y2 (blue), NG5 (red), Y2 + NG5 (green) HA proteins, or mock (PBS) (black). Pre-immune mice were vaccinated with mRNA LNPs encoding Y2 (orange), NG5 (navy blue), Y2 + NG5 (purple) HA proteins, or mock (PBS) (pink). The Y-axis denotes the Log 2 HAI titer of each individual mouse, and the X-axis indicates the virus tested. Dashed lines denote HAI titers of 1:40 (lower) and 1:80 (upper). Statistical significance was determined using a Kruskal-Wallis one-way ANOVA with Dunn’s multiple comparison test (*=p <0.05, **=p <0.01, ***=p <0.001, ****=p <0.0001).

Sera collected after the second vaccination were pooled for each vaccine group and evaluated for neutralization activity against panels of H1N1 and H3N2 viruses isolated between 2017 and 2023 (**Fig. 4**). Both influenza naïve and pre-immune mice immunized with vaccines containing Y2 HA exhibited strong neutralization activity against H1N1 viruses isolated from 2018 to 2022. On average, 50% neutralizing antibody titers in these groups were ∼14-18-fold higher than those observed in the mice immunized with monovalent NG5 HA or mock vaccines, which exhibited consistently low neutralizing activity against the H1N1 virus panel regardless of pre-immune status (**Fig. 4A**). For H3N2 viruses, influenza-naive and pre-immune mice immunized with NG5 HA had comparable neuralization activity across isolates from 2017 to 2023. In general, pre-immune mice vaccinated with NG5 HA mRNA LNPs had ∼2-fold higher 50% neutralization titers than naïve mice receiving the same vaccines. Notably, against the KS/17 isolate, pre-immune mice vaccinated with NG5 HA had ∼10-fold higher neutralization titers compared with influenza-naïve NG5 vaccinated mice. Additionally, influenza-naïve mice vaccinated with NG5 HA containing vaccines had ∼2-4-fold higher neutralization titers against the DC/23 strain than mice immunized with Y2 mRNA LNPs or mock vaccines, while pre-immune NG5 HA vaccinated mice had ∼4-5 fold higher neutralization titers relative to these groups. Mice immunized with Y2 HA mRNA LNP or mock vaccines had similar neutralization profiles against the H3N2 influenza virus panel regardless of pre-immune status, with 50% neutralization titers that were generally ∼8-14-fold lower than titers observed in NG5 HA mRNA vaccinated mice against H3N2 influenza viruses isolated between 2017 to 2020 (**Fig. 4B**).

**Figure 4.**
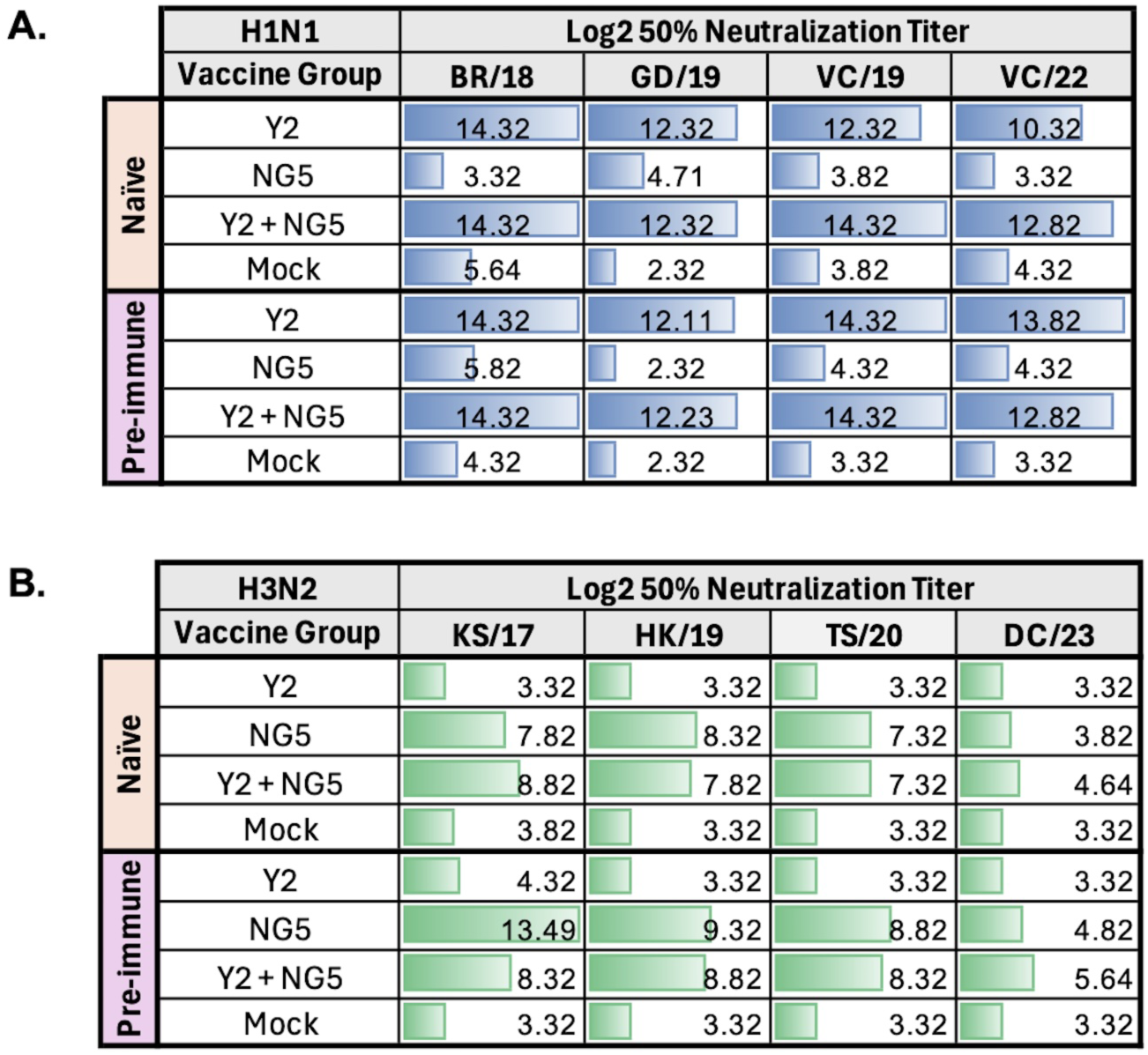
Day 42 Neutralizing antibody response. Serum samples collected from influenza naïve pre-immune mice two weeks after the second vaccination, day 42, were pooled for each group and examined for neutralization activity against historical H1N1 (A) and H3N2 (B) vaccine strains. The Log2 50% neutralization titer values are indicated in each cell, the vaccine groups are listed in the left most column, and the viruses assayed against are listed across the top row of the table. The colored bars in each cell correspond to the amount of neutralization observed ranging from 3.32 - 12.32. All assays were performed in duplicate and their results averaged. A Log2 titer of 3.32 represents the lower limit of detection (serum dilution of 1:10) and a Log2 titer of 12.32 represents the upper limit of detection (serum dilution of 1:5120) of the microneutralization assay.

A separate cohort of male (n=10) and female (n=10) mice was vaccinated in a prime boost regimen with a bivalent mixture of mRNA LNPs encoding COBRA Y2 and NG5 HA antigens to assess any potential sex-based differences in vaccine immunogenicity. Following two vaccinations, female mice developed significantly higher Y2 H1-specific IgG titers than males, whereas no significant differences were observed between sexes in NG5 H3-specific IgG responses (**Fig. 5A, B**). Both sexes had sero-protective HAI titers against H1N1 and H3N2 viruses isolated between 2017 and 2022. Although female mice exhibited significantly higher HAI titers against the BR/18 and VC/22 H1N1 strains (p < 0.05), and male mice had higher titers against the DR/21 H3N2 strain (p < 0.01), no significant sex-based differences were observed for the remaining viruses in either panel (**Fig. 5C, D**).

**Figure 5.**
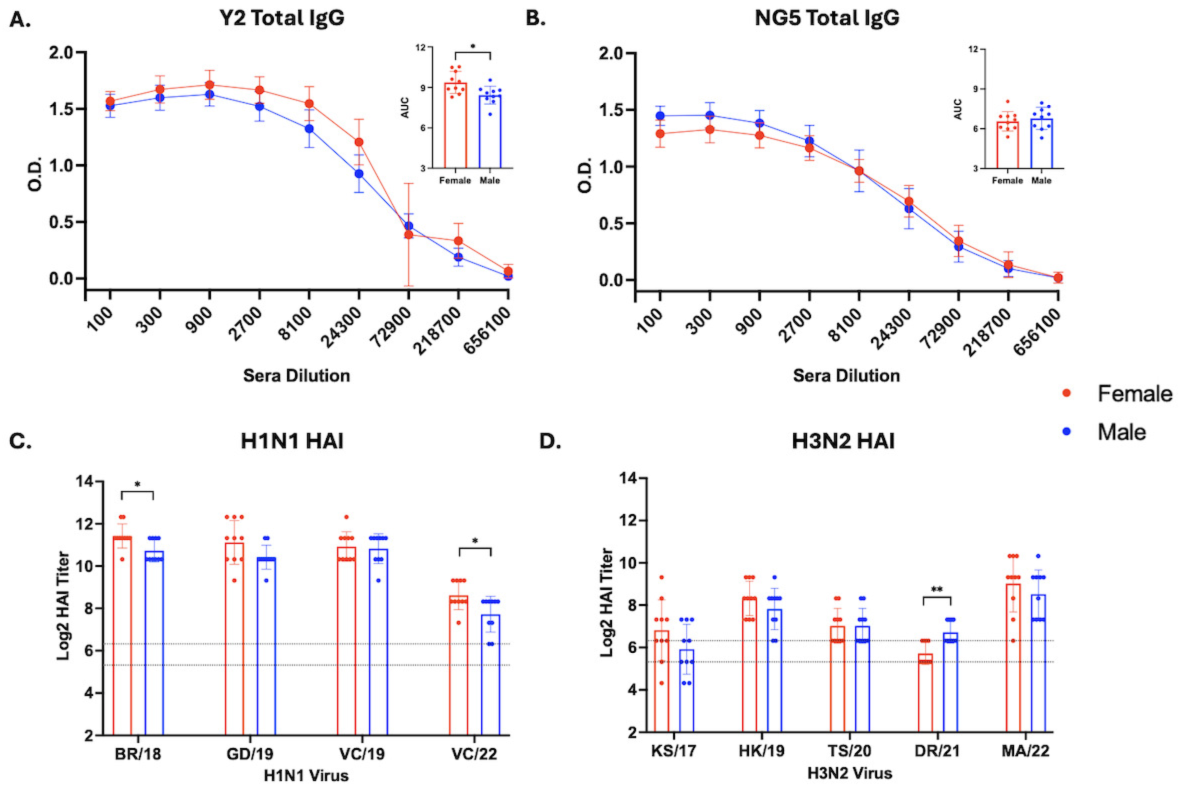
Comparison of humoral immune responses in male and female mice following COBRA mRNA LNP vaccination. Serum samples were collected from male (blue) and female (red) mice two weeks after the second vaccination, day 42, and evaluated for total IgG binding Y2 (A) and NG5 (B) rHA proteins by ELISA. Mean O.D. ± SEM is plotted as a function of serum dilution, and the AUC for each individual animal is shown in the corresponding panels. HAI antibody responses were also assessed against panels of H1N1 (C) and H3N2 (D) viruses. Statistical significance was determined using a Mann–Whitney U test (*=p <0.05, **=p <0.01, ***=p <0.001, ****=p <0.0001).

### COBRA mRNA LNP Vaccines elicit subtype specific antibody- and cytokine-secreting cells

Spleens were collected from a subset of mice (n=3/group) on day 37 following the second vaccination, pooled for each vaccine group, and evaluated for antigen-specific antibody secreting cells (ASCs) (**Fig. 6**). In general, vaccines encoding the Y2 H1 HA induced robust H1-specific responses, yielding ∼200-800 Y2 HA-specific ASCs and ∼150-650 BR/18-specific ASCs per million splenocytes. (**Fig. 6A, B**). On average, these mice also had ∼40-112 group 1 stalk-specific cH6/1 HA ASCs per million splenocytes (**Fig. 6C**). Pre-immune mice vaccinated with Y2 + NG5 mRNA LNPs had ∼2-3-fold higher frequencies of Y2, BR/18, and cH6/1 HA specific ASCs compared with naïve mice receiving the same vaccine. In contrast, mice immunized with the NG5 HA encoding mRNA or mock vaccines generated less than ∼30 H1 HA-specific ASCs per million splenocytes across the group 1 HA antigens (**Fig. 6A-C**). However, vaccines encoding the NG5 H3 HA elicited strong H3 HA-specific responses, with ∼350-1100 NG5 HA-specific ASCs and ∼55-180 group 2 stalk-specific cH7/3 HA ASCs per million splenocytes (**Fig. 6D, E**). As observed in the H1 HA-specific responses, pre-immune mice vaccinated with Y2 + NG5 HA mRNA LNPs had a ∼2-3-fold higher frequency of H3 HA-specific ASCs compared to naïve mice immunized with the same vaccine formulation. Mice vaccinated with Y2 HA encoding mRNA or mock vaccines had less than ∼12 H3 HA-specific ASCs per million splenocytes across the group 2 HA antigens (**Fig. 6D, E**).

**Figure 6.**
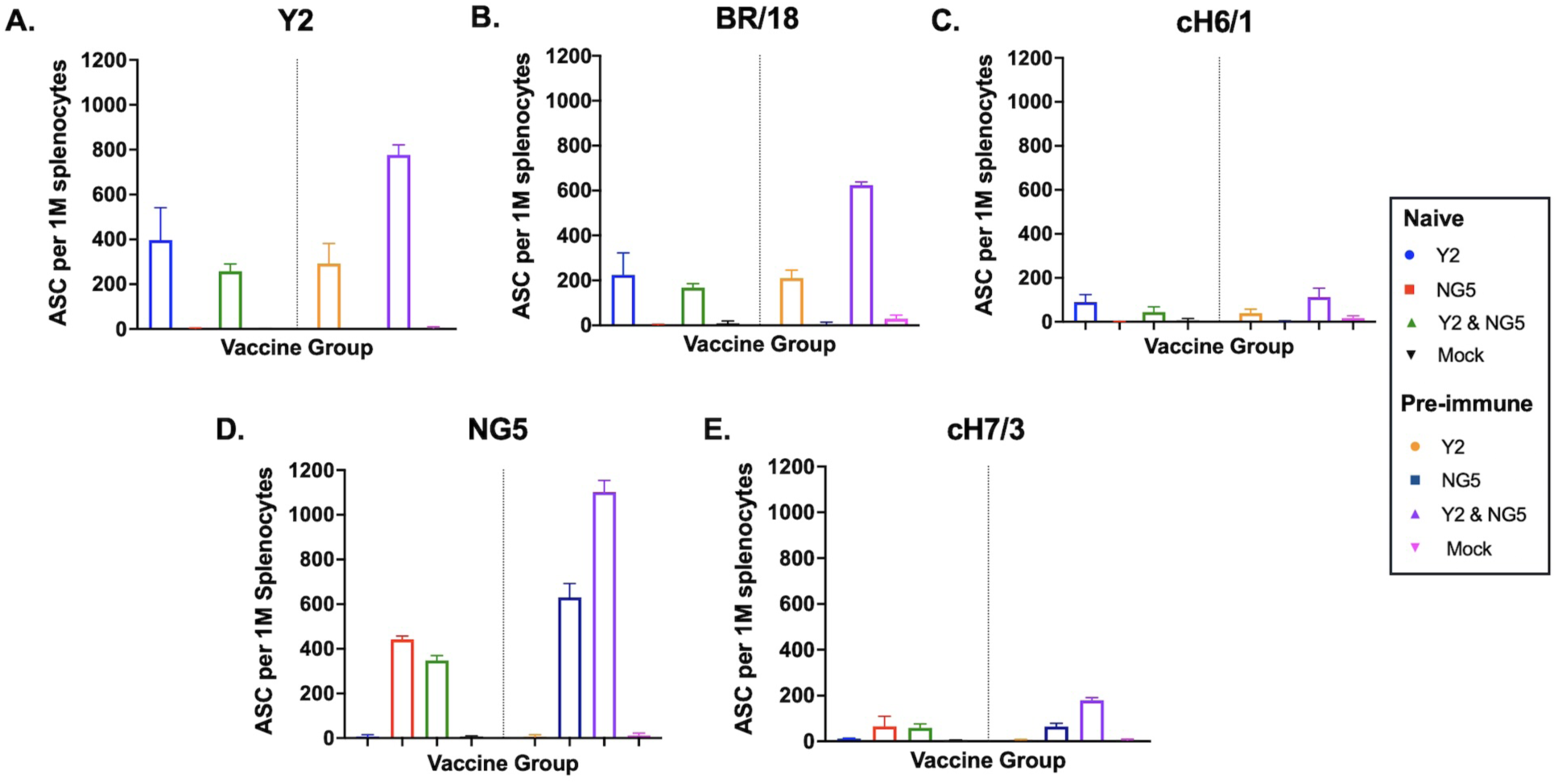
Frequency of antigen-specific antibody secreting cells. Splenocytes were collected from three mice per group following the second vaccination, day 37, pooled by vaccine group, and probed by ELISpot for reactivity against Y2 (A), BR/18 (B), cH6/1 (C), NG5 (D), and cH7/3 (E) rHAs. Bars represent the mean number of antibody secreting cells (ASC) per million splenocytes ± SD from triplicate wells. Naïve mice vaccinated with mRNA LNPs encoding Y2 (blue), NG5 (red), Y2 + NG5 (green) HA proteins, or mock (PBS) (black), and pre-immune mice vaccinated with mRNA LNPs encoding Y2 (orange), NG5 (navy blue), Y2 + NG5 (purple) HA proteins, or mock (PBS) (pink).

Pooled splenocytes from each group were also evaluated for H1 and H3 HA head (H) and stalk (S) specific interferon-gamma (IFN-γ) secreting cells (**Fig. 7**). Mice immunized with mRNA encoding the Y2 HA exhibited increased IFN-γ responses following stimulation with the H1 HA stalk peptide pool. On average, pre-immune mice displayed ∼6.5-fold higher frequencies of H1 HA stalk-specific IFN-γ secreting cells compared with naïve mice, and a more modest, ∼1.3-fold, increase in H1 HA head-specific IFN-γ responses (**Fig. 7A**). Vaccination with NG5 HA encoding mRNA preferentially elicited IFN-γ secreting cells specific to the H3 HA stalk peptide pool, with an ∼3.5-fold higher frequency than those stimulated by the H3 HA head peptide pool. Pre-immune mice immunized with vaccines encoding NG5 had ∼2.6-fold and ∼1.4-fold higher frequencies of H3 HA head- and stalk-specific IFN-γ secreting cells than naïve mice (**Fig. 7B**). Mice immunized with the mock vaccine or those encoding Y2 HA had minimal H3 HA-specific IFN-γ, with fewer than ∼45 cytokine secreting cells per million splenocytes (**Fig. 7B**).

**Figure 7.**
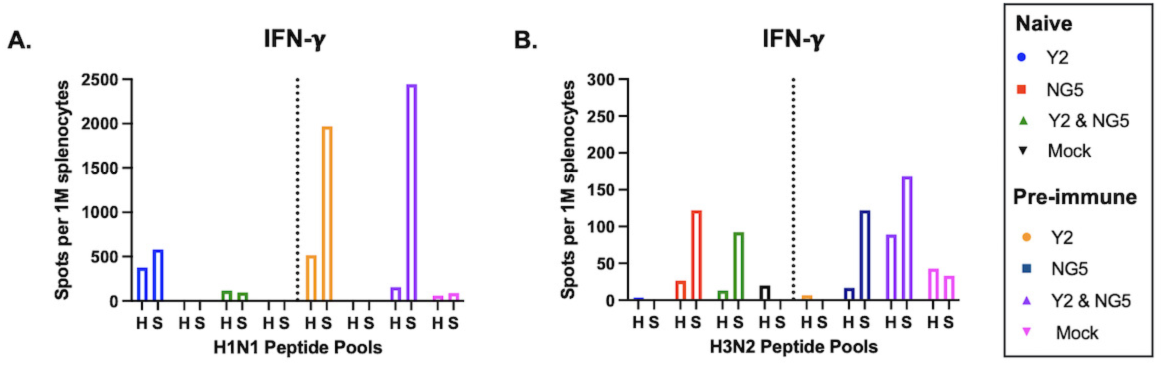
Frequency of antigen-specific cytokine secreting cells. Splenocytes were harvested from three mice in each group following the second vaccination on day 37. The cells were pooled for each group and examined by ELISpot for IFN-γ secretion upon exposure to H1N1 (A) and H3N2 (B) HA head (H) and stalk (S) peptide pools. Cytokine secreting cell were enumerated as individual spots, and the number of spot forming cells per million splenocytes is plotted on the Y-axis. Naïve mice were vaccinated with mRNA LNPs encoding Y2 (blue), NG5 (red), Y2 + NG5 (green) HA proteins, or mock (PBS) (black), and pre-immune mice vaccinated with mRNA LNPs encoding Y2 (orange), NG5 (navy blue), Y2 + NG5 (purple) HA proteins, or mock (PBS) (pink). Bars represent the mean response for each vaccine group.

### Y2 COBRA mRNA vaccines protected mice from lethal H1N1 infection

Following the second vaccination, on day 56, mice were intranasally challenged with a lethal dose of the BR/18 H1N1 virus. All influenza naïve and pre-immune mice vaccinated with mRNA LNPs encoding the Y2 COBRA HA antigen survived the challenge, with less than 5% weight loss and no detectable clinical signs of infection up to fourteen days post challenge (**Fig. 8A-C**). In comparison, both influenza naïve and pre-immune mice vaccinated with monovalent NG5 HA mRNA LNPs or mock vaccines had influenza virus associated clinical signs as early days 2-3 post infection, with ∼20% body weight loss, and reached humane endpoint criteria requiring euthanasia by days 5-6 post infection (**Fig. 8A-C**). Consistent with these outcomes, mice immunized with Y2 encoding mRNA vaccines had no detectable viral loads in the lungs at day 3 post infection, whereas mice vaccinated with monovalent NG5 HA mRNA LNPs or mock vaccines had <1×10^6^ PFU of BR/18 virus in the lungs at the same time point (**Fig. 8D**).

**Figure 8.**
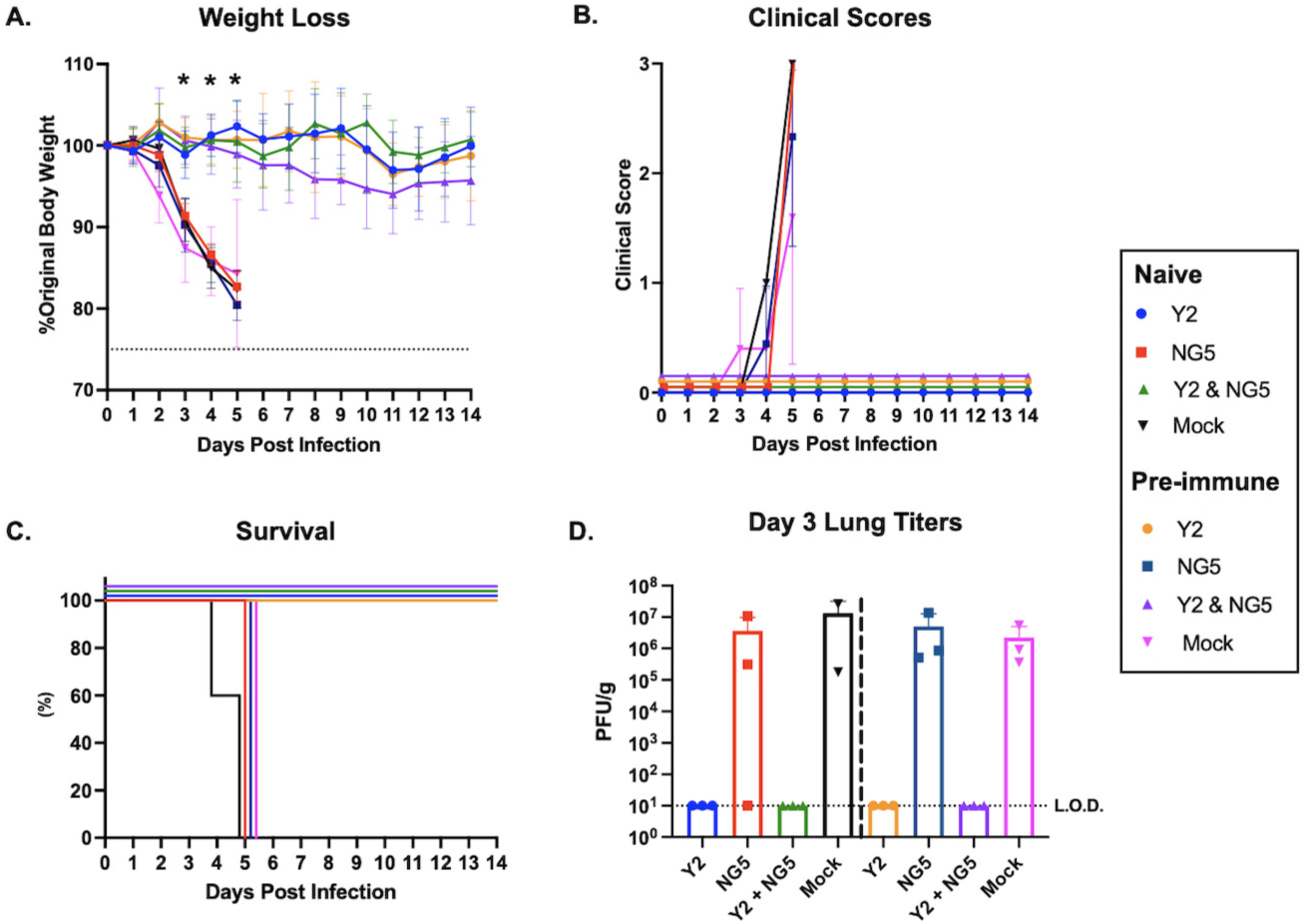
Protective efficacy of COBRA mRNA LNP vaccines against BR/18 H1N1 challenge in naïve and pre-immune mice. On day 56, remaining mice were challenged intranasally with BR/18 H1N1 virus, and monitored for weight loss (A), clinical disease scores (B), and survival (C) for 14 days. Influenza naïve mice were vaccinated with mRNA LNPs encoding Y2 (blue), NG5 (red), Y2 + NG5 (green) HA proteins, or mock (PBS) (black), and pre-immune mice (D-F) were immunized with mRNA LNPs encoding Y2 (orange), NG5 (navy blue), Y2 + NG5 (purple) HA proteins, or mock (PBS) (pink). Lungs samples collected three days post-infection were assessed by plaque assay to quantify viral burden (D). Viral titers are expressed as plaque forming units (PFU) per gram of lung tissue (PFU/g). The dotted line indicates the limit of detection (LOD). Statistical significance was determined using a Kruskal-Wallis one-way ANOVA with Dunn’s multiple comparison test (*=p <0.05, **=p <0.01, ***=p <0.001, ****=p <0.0001).

Similarly, male and female mice vaccinated with Y2 and NG5 mRNA-LNP vaccines were completely protected from BR/18 H1N1 challenge. No significant differences in weight loss were observed between sexes, and neither group lost more than 5% of body weight following infection. Furthermore, vaccinated animals exhibited no clinical signs of disease and had no detectable virus in the lungs on day 3 post-infection. In contrast, mock-vaccinated mice (n = 10) developed clinical signs beginning on day 2 post-infection and lost significantly more weight than vaccinated animals starting on day 2 (*p* < 0.05), with peak losses approaching 25% of body weight. Mock-vaccinated animals also had significantly higher lung viral titers than the vaccinated mice (p < 0.0001) reaching ∼1 × 10⁶ PFU/g on day 3 post-infection, and all succumbed to infection by day 7 (**Fig. 9A-D**).

**Figure 9.**
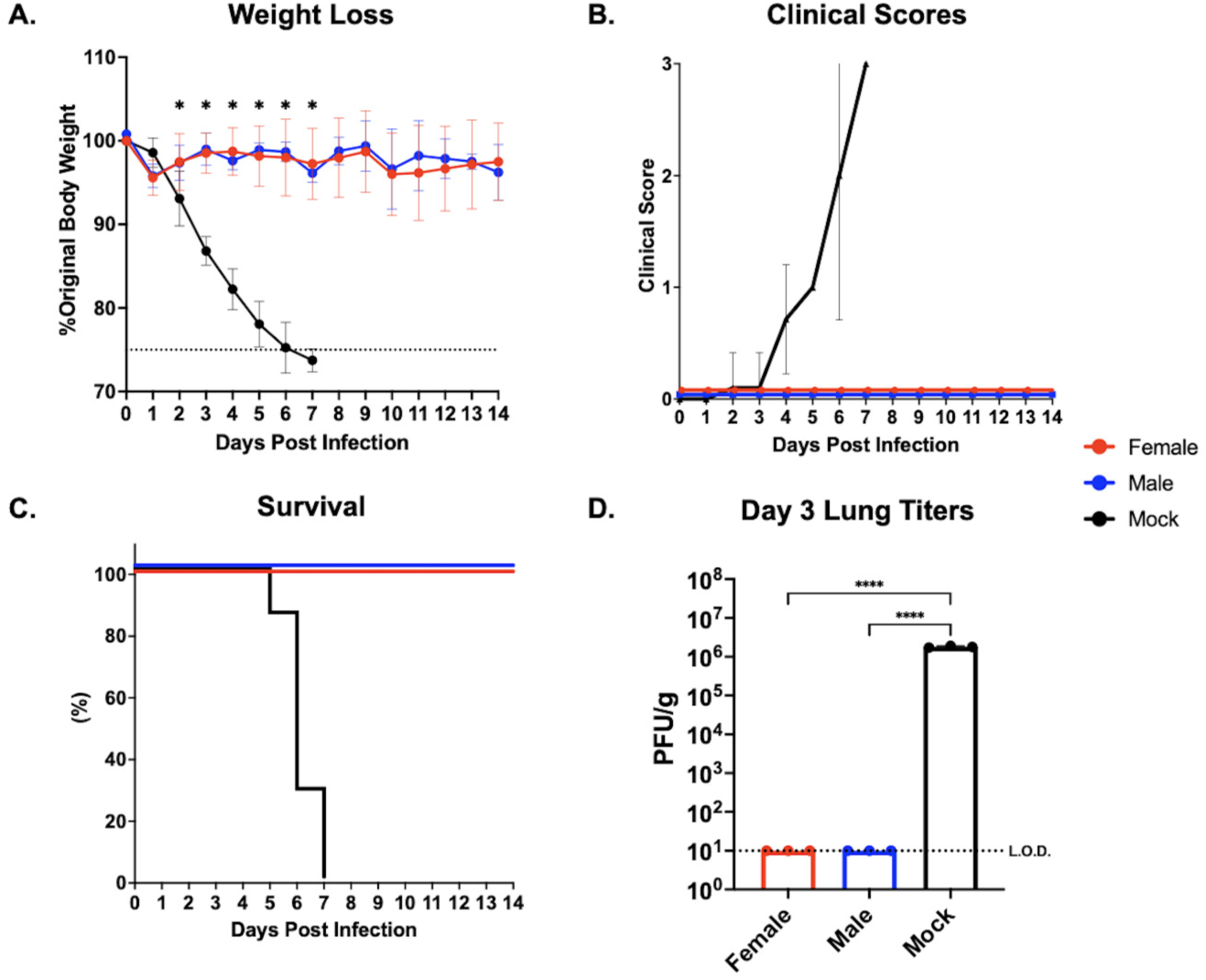
Protective efficacy of COBRA mRNA LNP vaccines against BR/18 H1N1 challenge in male and female mice. Male (blue; n = 10), female (red; n = 10) mice vaccinated with Y2 and NG5 mRNA LNP vaccines, along with mock vaccinated mice (black; n = 10) were challenged intranasally with BR/18 H1N1 virus, and monitored for weight loss (A), clinical disease scores (B), and survival (C) for 14 days. Lungs samples collected three days post-infection were assessed by plaque assay to quantify viral burden (D). Viral titers are expressed as plaque forming units (PFU) per gram of lung tissue (PFU/g). The dotted line indicates the limit of detection (LOD). Statistical significance was determined using a Kruskal-Wallis one-way ANOVA with Dunn’s multiple comparison test (*=p <0.05, **=p <0.01, ***=p <0.001, ****=p <0.0001).

## Discussion

The COBRA HA mRNA LNP vaccines elicited broadly protective, subtype-specific immune responses in both influenza naïve and pre-immune animals, preventing weight loss and suppressing viral replication in the respiratory tract. From a humoral perspective, COBRA H1 and H3 mRNA vaccines generated robust antigen-specific IgG, HAI, and neutralizing antibody responses against historical and contemporary influenza strains. This breadth across temporal antigenic space supports the COBRA HA design strategy as an effective approach to mitigate the effects of antigenic drift and seasonal vaccine mismatch. While H1N1 HAI titers were comparable between naïve and pre-immune mice, H3N2 HAI titers tended to be higher in pre-immune animals, although these differences were generally not statistically significant. Female mice also developed higher total H1-specific IgG responses than males. However, differences in HAI activity were limited, with females exhibiting significantly higher titers against two of the four H1N1 viruses and males exhibiting higher titers against a single H3N2 strain. No significant differences were observed across the remaining viruses, H3-specific IgG responses, or protection from viral challenge. Together, these findings suggest that COBRA mRNA vaccination elicits broadly comparable immune responses across sexes, contrasting with reports that females often mount consistently stronger humoral responses following vaccination with egg-based inactivated influenza vaccines (47–49). Lastly, combining H1 and H3 COBRA HA antigens in a bivalent formulation did not significantly reduce their immunogenicity, supporting the feasibility of future multivalent vaccine strategies analogous to current trivalent seasonal influenza vaccines.

Beyond humoral immunity, COBRA HA mRNA vaccination elicited robust cellular responses characterized by the expansion of antigen-specific antibody and cytokine secreting cells. ASC responses were primarily directed toward the HA head and were more frequent in pre-immune animals, consistent with the immunodominance of HA head-specific antibody responses typically elicited by inactivated influenza vaccines (50). In contrast, cytokine-secreting cells preferentially recognized HA stalk-derived peptides, indicating recruitment of cellular immune responses directed against conserved regions that may support cross-strain protection (51, 52). Pre-existing immunity further amplified these responses, likely through reactivation of memory cell populations targeting conserved HA epitopes shared amongst the COBRA HA antigens and the previously encountered strains (53). This was reflected in increased frequencies of IFN-γ-secreting cells in pre-immune animals. The predominance of IFN-γ responses in this study is consistent with the Th1-biased immunity commonly elicited by mRNA-LNP vaccines (54). IFN-γ production by Th1 cells and cytotoxic T lymphocytes is associated with enhanced viral clearance and improved disease outcomes (55). Mechanistically, the mRNA LNP vaccines likely promote innate immune activation through their ionizable lipid components, driving pro-inflammatory cytokine production and enhancing antigen presentation, thereby supporting robust adaptive immune priming (28, 54).

These findings are highly relevant to human populations, where pre-existing influenza immunity is nearly universal (56, 57). The ability of COBRA HA mRNA vaccines to leverage immunological memory while expanding responses to conserved HA epitopes highlights their potential to improve cross-protective efficacy against antigenically drifted influenza strains. mRNA HA vaccine platforms are well suited for next-generation influenza vaccine strategies due to their adaptability and ability to be rapidly manufactured (58). This aligns synergistically with the COBRA design approach, enabling timely updates of optimized antigens based on the evolving landscape of circulating viral strains. Unlike egg-based systems, mRNA vaccines can be produced at scale within shorter timeframes and avoid propagation-associated mutations and species-specific glycosylation biases by driving in situ protein expression in host cells (29, 59, 60). Current clinical trials evaluating wild-type HA based mRNA vaccines demonstrate the promise of this platform, however these approaches typically elicit strain specific immune responses with limited breadth against antigenically drifted variants (7). Incorporating broadly reactive COBRA antigens into mRNA LNP delivery platforms has the potential to overcome these limitations by expanding antigenic coverage across diverse and emerging strains.

Despite these encouraging findings, the limitations of this study should be acknowledged. First, the pre-immune models relied on exposure to a single H1N1 and H3N2 virus and therefore do not fully recapitulate the complex and evolving influenza exposure histories observed in humans. Nevertheless, these models demonstrate that COBRA mRNA vaccines can effectively leverage pre-existing immunity to enhance both humoral and cellular responses, suggesting that vaccine performance may be further strengthened in individuals with more extensive immune histories. Second, although broad reactivity was observed across the H1N1 virus panel, H3N2-specific responses were not universally protective. The rapid antigenic evolution of H3N2 viruses presents a greater challenge for achieving consistent coverage, highlighting opportunities to further refine COBRA H3 HA antigen design or explore multivalent H3 formulations. Finally, the durability of COBRA mRNA-LNP-induced immunity was not evaluated. Because long-term protection remains a critical consideration for mRNA-based vaccines, future studies should assess the longevity of both humoral and cellular responses and determine the duration of protection following vaccination (61, 62).

Overall, these findings support the integration of COBRA HA antigen design with mRNA vaccine platforms as a scalable strategy to improve seasonal influenza vaccine efficacy by expanding immune breadth against both contemporary and future influenza strains. COBRA HA mRNA vaccines elicited functional broadly reactive HAI and neutralizing antibodies, while also stimulating robust antibody and cytokine secreting effector cell populations. Notably, the enhanced responses observed in pre-immune animals demonstrate the ability of these vaccines to effectively engage immune memory to elicit robust immune responses. Together, the broad protection observed across multiple influenza subtypes and immune backgrounds highlight the translational potential of COBRA mRNA vaccines and supports their continued development as next-generation influenza vaccines capable of addressing the challenges posed by ongoing viral evolution.

## Materials and Methods

### Vaccine Antigens

The H1 COBRA HA, Y2, was designed using wild-type H1 HA sequences from viruses that circulated during May 2014 – September 2016 as previously described (45). The H3 COBRA HA, NG5, was synthesized from wild-type H3 HA sequences from viruses that circulated during October 2018 to April 2021 as previously described (46). The nucleotide sequences for each COBRA HA were optimized for messenger ribonucleic acid (mRNA) expression and individually cloned into separate mRNA expression vectors (TriLink Biotechnologies, San Diego, CA, USA) and encapsulated in lipid nanoparticles (LNPs) (Acuitas, Vancouver, BC, Canada).

### Viruses

Influenza viruses were obtained from Virapur (San Diego, CA), the International Reagent Resources (IRR), the Biodefence and Emerging Infections Research Resource (BEI Resources), or the United States Center for Disease Control and Prevention (CDC). All viruses were passaged once in embryonated chicken eggs following World Health Organization (WHO) established protocols (63). After propagation and harvest, viruses were stored at −80 °C in individual 1 mL aliquots. H1N1 viruses: A/California/07/2009 (CA/09), A/Michigan/45/2015 (MI/15), A/Brisbane/02/2018 (BR/18), A/Guangdong-Maonan/SWL1536/2019 (GD/19), A/Victoria/2570/2019 (VC/19), and A/Victoria/4897/2022 (VC/22). H3N2 viruses: A/Switzerland/9715293/2013 (SW/13), A/Kansas/14/2017 (KS/17), A/South Australia/34/2019 (SA/19), A/Hong Kong/2671/2019 (HK/19), A/Tasmania/503/2020 (TS/20), A/Darwin/9/2021 (DR/21), A/Massachusetts/18/2022 (MA/22), and A/District of Columbia/27/2023 (DC/23). One H1N1 virus, A/Singapore/6/1986 (SG/86) and one H3N2 virus, A/Panama/2007/1999 (PN/99) were used to establish an immune history to seasonal influenza in a cohort of mice and ferrets.

### Vaccination of Mice

A cohort of female DBA/2J mice (n=112), *Mus musculus*, 6-8-weeks old, were purchased from The Jackson Laboratory (Bar Harbor, ME). At the beginning of the study, mice were randomly divided into groups (n=15 mice/group (experimental vaccine groups) n=11 mice/group (mock vaccinated control groups)). On day −28, half of the mice (n=56) were weighed and infected with 50 μL of a mixture containing 5×10^5^ plaque forming units (PFU) of SG/86 H1N1 virus and 5×10^5^ PFU of PN/99 H3N2 virus diluted in sterile phosphate buffered saline (PBS) (Thermo Fisher, Waltham, MA) to establish an immune history to historical influenza viruses. Following infection, the mice were monitored for weight loss and clinical symptoms associated with influenza infection daily for 14 consecutive days. The mice were then allowed to rest for 14 additional days prior to vaccination. On day 0, the previously immunized (pre-immune) mice (n=15/group) and a set of influenza naïve mice (n=15/group) were vaccinated intramuscularly in the hind leg with 1 μg of Y2, 1 μg of NG5, or 1μg of Y2 and 1 μg of NG5 mRNA LNPs diluted in PBS to a total volume of 50 μL. Separate groups of pre-immune and naïve mice (n=11/group) were vaccinated intramuscularly with 50 μL of PBS to serve as mock vaccinated controls. Two weeks after vaccination blood was collected from each mouse on day 14 to assess influenza specific antibody responses elicited by the vaccines. On day 28 each mouse was vaccinated intramuscularly with homologous vaccines to those that were administered on day 0. Nine days later, on day 37, three mice from each group were humanely euthanized. Spleens from these animals were harvested and homogenized into single cell suspensions to analyze influenza specific cellular immune responses. Two weeks after the second vaccination, on day 42, blood was collected from each mouse, and on day 56 the mice were intranasally infected with 4×10^4^ PFU of BR/18 H1N1 influenza virus that was diluted in PBS to a total volume of 50 μL. Following infection, the mice were monitored for weight loss and clinical symptoms associated with influenza infection daily for 14 consecutive days. On day 59 and day 62, lungs were collected from three mice in each group to assess for the presence of influenza virus. On day 70 the mice were humanely euthanized and the study was terminated.

A separate cohort of influenza-naïve DBA/2J mice (10 females and 10 males; 6–8 weeks old; The Jackson Laboratory) was acclimated for 72 h and vaccinated intramuscularly on days 0 and 28 with 1 μg each of Y2 and NG5 mRNA-LNP vaccines. An additional group of age matched female DBA/2J mice (n = 10) were vaccinated with PBS and served as mock-vaccinated controls. Blood samples were collected on days 14 and 42 to assess vaccine-induced antibody responses. On day 56, mice were challenged intranasally with 4 × 10^4 PFU of BR/18 H1N1 virus. Following challenge, animals were monitored daily for 14 days for weight loss and clinical signs of disease. Lung samples were collected from a subset of mice (n = 3/group) on day 59 to quantify viral replication, and the study was terminated on day 70.

All mice were housed in microisolator units and provided unlimited access to food and water for the totality of the study. Each animal was cared for under the National Institutes of Health guide for the care and use of laboratory animals, and all procedures were approved by the Cleveland Clinic Institutional Animal Care and Use Committee (IACUC) (protocol #2935) prior to implementation.

### Hemagglutination Inhibition Assay

The hemagglutination inhibition assay (HAI) was performed as previously described (64). HAIs involving H1N1 viruses were performed with 0.8% turkey red blood cells (RBCs) and HAIs involving H3N2 isolates were conducted with 0.75% guinea pig RBCs (Lampire Biological Laboratories, Pipersville, PA, USA) supplemented with 20 nM Oseltamivir Carboxylate (Aobious, Gloucester, MA, USA). The HAI titer of the antibodies in the serum was calculated using the reciprocal of the dilution corresponding to the last well that had non-agglutinated RBCs. For this study, a “sero-protective” HAI antibody titer was defined as a titer ≥ 1:40, as per the WHO guidelines for evaluating influenza vaccines.(65)

### Enzyme-Linked Immunosorbent Assay

Enzyme-Linked Immunosorbent Assays (ELISAs) were performed as previously described (66). Individual 96-well plates were coated with 1 μg/mL of either Y2, BR/18, a chimeric HA with the H6 head of A/Mallard/Sweden/81/2002 and the H1 stalk from A/California/07/2009 (cH6/1), NG5, TS/20, or a group 2 chimeric HA with the H7 head of A/Anhui/1/2013 and the H3 stalk from A/Texas/50/2012 (cH7/3) (67) rHA in carbonate coating buffer (pH 9.4). Next,100 μL of pooled sera samples from each group were diluted 1:100 in blocking buffer and added to the first column of each plate and serially diluted 2-fold across the plate. After washing, 100 μL of Goat anti-mouse IgG HRP (1 mg/mL) (Cat. no. 1030-05) (Southern Biotech, Birmingham, AL, USA) secondary antibody diluted 1:4000 in blocking buffer was added to each well and incubated at 37° C for 90 min. Plates were then washed, and 100 μL of substrate (1 mg/mL 2,2’-azino-bis(3-ethylbenzothiazoline-6-sulfonic acid (ABTS) diammonium salt (Sigma, Burlington, MA, USA), resuspended in McIlvaine’s solution (pH 5.0) was added to each well and allowed to develop for 12-15 min at 37° C in a humidified chamber. Next, 50 μL of a 1% sodium dodecyl sulfate (SDS) (Thermo Fisher, Waltham, MA, USA) solution was added to each well to stop the colorimetric reaction. The plates were read at 414 nm using a BioTek Epoch 2 plate reader (Agilent, Santa Clara, CA, USA) equipped with Gen6 software. Endpoint dilutions were determined by taking the reciprocal dilution of the most dilute well that contained greater than twice the mean signal of the background wells.

### Neutralization Assay

Neutralization assays were performed as previously described with some modifications (46). In brief, pooled sera samples were treated with receptor destroying enzyme (RDE) (Denka Seiken Co., Tokyo, Japan) according to the manufacturer’s instructions. Sera samples were diluted two-fold across the plate starting from a 1:10 dilution and allowed to incubate with 50 μL of 100TCID50 of virus for 1 hour. Next, 100 μL of Madin Darby Canine Kidney (MDCK) cells (BEI Resources) (1.5×10^5^ cells) were added to each well and the plates were incubated for 16-20 hours at 37 °C, 5% CO2. The last column of the plate was reserved for virus only and cell only control wells. Plates were then washed, fixed by adding 100 μL of acetone (4 °C), and allowed to air dry for 10-12 hours. The plates were then washed three times and probed with 100 μL/well of rabbit pan influenza A nucleoprotein primary antibody (Sino Biological, Houston, TX, USA, RRID# AB_3677055, cat# 40208-R061) diluted 1:10,000 in wash buffer containing 5% non-fat dry milk for 1 hour at room temperature. The plates were then washed three times and probed with 100 μL/well of goat anti-rabbit IgG-Fc horseradish peroxidase (HRP) secondary antibody (Sino Biological, RRID# AB_2814815, cat# SSA003) diluted 1:20,000 000 in wash buffer containing 5% non-fat dry milk for 1 hour at room temperature. The plates were subsequently washed three times and 100 μL/well of substrate (phosphate citrate buffer (Sigma), o-phenylenediamine dihydrochloride (OPD) tablets (Sigma), 0.05%, 30% H2O2) was added to each well. The plates were allowed to develop for 10-20 mins and 100 μL of stop solution (2N H2SO4 (Thermo Fisher) was added to each well. Plates were read at 490nm a BioTek Epoch 2 plate reader (Agilent) using Gen6 software. The 50% neutralization titer was determined by comparing the optical density of each well to that of the virus only control wells after subtracting the average signal from the cell only control wells. All samples were run in duplicate, and the results were averaged across replicates.

### Cytokine Secreting Cell ELISpot

Cytokine secreting cell Enzyme-linked Immunospot (ELISpot) assays were performed as previously described with some modifications (46). Individual 14 and 15-mer peptides, overlapping by 7 amino acids, spanning the entire HA sequence of A/New York/18/2009 (H1N1) (NY/04) (Catalog No. NR-19245) and A/Perth/16/2009 (H3N2) (PE/09) (Catalog No. NR-19266) were obtained from BEI Resources and separated into peptide pools comprising the HA head and stalk regions at a final concentration of 2ug/mL per peptide (68). Mouse IFN-γ Single-Color ELISpot Kits (Cellular Technology Limited (CTL), Shaker Heights OH, USA cat# mIFNg-1M) were used to enumerate IFN-γ secreting cells according to the manufacturer’s instructions (69). Splenocytes were pooled from each group of mice and added to each well (3 x 10^5^ cells/well) containing peptide pools. Data were acquired using the CTL ImmunoSpot^®^ reader (CTL) and analyzed with ImmunoSpot^®^ Version 7 software. Each spot corresponds to an individual cytokine-secreting cell. Background responses from negative control wells were subtracted to normalize the data across all experimental groups.

### Antibody Secreting Cell ELISpot

Splenocyte samples were pooled for each group of mice and assessed for antibody secreting cells (ASCs) specific to rHA antigens as previously described (70). One day prior to the assay, MultiScreenHTS 96-well clear plates with PVDF membrane filters (Sigma) were coated with 1.5- 2.5 μg of either Y2, BR/18, cH6/1, NG5, or cH7/3 rHA resuspended in 50 μL PBS. Pooled splenocytes were added to the plates in 3-fold serial dilutions starting at 300,000 cells/well and incubated at 37 °C, 5% CO2 for 16-18h. Next, 50 μL of a solution containing Goat anti-Mouse IgG, Human ads-AP (Southern Biotech, Birmingham, AL, USA cat. no. 1030-04) diluted 1:4000 was added to each well and plates were incubated for 2 h at 37°C. After incubation, plates were washed and 50 μL of Nitro-Blue Tetrazolium and 5-bromo-4-chloro-3’-indolyphosphate (NBT/BCIP) substrate (Thermo Fisher) was added to each well. The substrate was allowed to develop for 10-15 min at 37°C until visible spots clearly formed in each well. Spots were imaged using a CTL ImmunoSpot^®^ analyzer (CTL) and counted using ImmunoSpot^®^ software version 7.0.38.2 Professional SC.

### Influenza Plaque Assay

Influenza plaque assays were performed as previously described (42). In brief, 18-24h before the assay 1×10^6^ Madin-Darby Canine Kidney (MDCK) cells (Sigma) were added to each well of a six-well plate (Thermo Fisher) and incubated overnight at 37°C + 5% CO2. The next day, mouse lung tissues were weighed and homogenized in Dulbecco’s Modified Eagle Medium (DMEM) supplemented with 1% Penicillin/Streptomycin (DMEM + P/S) (Thermo Fisher). Lung homogenates were pelleted at 2,000 rpm for 5 min, the supernatants were harvested, and diluted in 10-fold serial dilutions in DMEM + P/S. The MDCK cells were washed and 100 μL of each dilution of the lung homogenates were added to the plates in duplicate wells. The plates were incubated for 1 h with agitation every 15 mins. Following incubation each well was washed twice, and a 50:50 v/v mixture of 2X Minimal Essential Medium (MEM),1.6% agarose, and 1 μg/mL of TPCK Trypsin (Thermo Fisher) was added to each well. The plates were incubated at 37°C + 5% CO2 for ∼72 h. After incubation, the solidified agarose gels were manually removed, and the cells were fixed for 10 min with 10% buffered formalin (Thermo Fisher). The plates were then stained with 1% crystal violet (Thermo Fisher) for 10 min, washed 5 times with fresh water, and allowed to air dry for 24 hours. Individual plaques were counted and reported as the number of plaques in the reciprocal of each dilution. The lung vial titers were presented as plaque forming units per gram of lung tissue (PFU/g) by dividing the number of viral plaques by the weight of each lung sample.

### Statistical Analysis

Statistical analyses were performed using GraphPad Prism (GraphPad Software, San Diego, CA, USA). Differences in ELISA, HAI, plaque assay, and weight-loss data among vaccine groups in naïve and pre-immune mice were evaluated using a Kruskal–Wallis one-way analysis of variance with Dunn’s multiple-comparison test. Differences in total IgG and HAI titers between male and female mice were assessed using a Mann–Whitney U test. Statistical significance was defined as p < 0.05 and deonted as follows (*=p <0.05, **=p <0.01, ***=p <0.001, ****=p <0.0001).

## Data Availability

The original data presented in this study are made publicly available via the National Institutes of Health (NIH) ImmPort online database. The data will also be made available to qualified members of the scientific research community upon written request.

## Acknowledgements

This project was funded as part of the Collaborative Influenza Vaccine Innovations Centers (CIVICs) by the National Institute of Allergy and Infectious Diseases (NIAID), a component of the NIH, Department of Health and Human Services, under contract 75N93019C00052. The funders had no role in study design, data collection and interpretation, or the decision to submit the work for publication. The authors would like to thank the Influenza Reagent Resource (IRR), Influenza Division, WHO Collaborating Center for Surveillance, Epidemiology, and Control of Influenza, Centers for Disease Control and Prevention (Atlanta, GA, USA) for providing some of the A(H1N1) and A(H3N2) influenza viruses. They would also like to thank the Cleveland Clinic Animal Resource staff, technicians, and veterinarians for their excellent animal care. Some figures were created with BioRender.com. JDA and TMR are inventors on patents PCT/US22/32799, PCT/US19/41679, PCT/US19/41704.

J.D.A. and T.M.R. conceptualized the experiments. J.M., M.T., and A.L. conducted the animal work and collected samples. C.C wrote the cytokine secreting cell ELISpot methods section. C.C., J.M., M.T., and A.L. performed serological assays and prepared the figures. J.D.A. analyzed the data and wrote the manuscript with input from T.M.R. All authors read and approved the final version of the manuscript.

## Figure Legend

**Supplemental Figure 1. Individual antigen-specific IgG endpoint titers following COBRA mRNA LNP vaccination.** Serum samples collected from individual mice two weeks after the second vaccination, day 42, were evaluated by ELISA for total IgG binding to Y2 (A) and NG5 (B) COBRA rHA proteins. Endpoint dilution titers for each animal are plotted on the Y-axis, and vaccine groups are listed on the X-axis. Statistical significance was determined using a Kruskal-Wallis one-way ANOVA with Dunn’s multiple comparison test (*=p <0.05, **=p <0.01, ***=p <0.001, ****=p <0.0001).

**Supplemental Figure 2. HAI antibody responses following a single COBRA mRNA LNP vaccination.** Serum samples collected from naïve (A, C) and pre-immune mice (B, D) two weeks after the first vaccination, day 14, were evaluated for HAI activity against historical H1N1 (A, B) and H3N2 (C, D) vaccine strains. Naïve mice were vaccinated with mRNA LNPs encoding Y2 (blue), NG5 (red), Y2 + NG5 (green) HA proteins, or mock (PBS) (black). Pre-immune mice were vaccinated with mRNA LNPs encoding Y2 (orange), NG5 (navy blue), Y2 + NG5 (purple) HA proteins, or mock (PBS) (pink). The Y-axis denotes the Log 2 HAI titer of each individual mouse, and the X-axis indicates the virus tested. Dashed lines denote HAI titers of 1:40 (lower) and 1:80 (upper). Statistical significance was determined using a Kruskal-Wallis one-way ANOVA with Dunn’s multiple comparison test (*=p <0.05, **=p <0.01, ***=p <0.001, ****=p <0.0001).

